# A chromosome-level assembly of the Atlantic herring – detection of a supergene and other signals of selection

**DOI:** 10.1101/668384

**Authors:** Mats E. Pettersson, Christina M. Rochus, Fan Han, Junfeng Chen, Jason Hill, Ola Wallerman, Guangyi Fan, Xiaoning Hong, Qiwu Xu, He Zhang, Shanshan Liu, Xin Liu, Leanne Haggerty, Toby Hunt, Fergal J. Martin, Paul Flicek, Ignas Bunikis, Arild Folkvord, Leif Andersson

## Abstract

The Atlantic herring is a model species for exploring the genetic basis for ecological adaptation, due to its huge population size and extremely low genetic differentiation at selectively neutral loci. However, such studies have so far been hampered because of a highly fragmented genome assembly. Here, we deliver a chromosome-level genome assembly based on a hybrid approach combining a *de novo* PacBio assembly with Hi-C-supported scaffolding. The assembly comprises 26 autosomes with sizes ranging from 12.4 to 33.1 Mb and a total size, in chromosomes, of 726 Mb. The development of a high-resolution linkage map confirmed the global chromosome organization and the linear order of genomic segments along the chromosomes. A comparison between the herring genome assembly with other high-quality assemblies from bony fishes revealed few interchromosomal but frequent intrachromosomal rearrangements. The improved assembly makes the analysis of previously intractable large-scale structural variation more feasible; allowing, for example, the detection of a 7.8 Mb inversion on chromosome 12 underlying ecological adaptation. This supergene shows strong genetic differentiation between populations from the northern and southern parts of the species distribution. The chromosome-based assembly also markedly improves the interpretation of previously detected signals of selection, allowing us to reveal hundreds of independent loci associated with ecological adaptation in the Atlantic herring.

## INTRODUCTION

The Atlantic herring (*Clupea harengus*) is a model system to study the genetic basis for ecological adaptation and the consequences of natural selection (Martinez Barrio *et al*., 2016; Lamichhaney *et al*., 2017). A major merit of this system for evolutionary studies is the minute genetic drift due to the enormous population size facilitating the detection of how natural selection affects the populations. The Atlantic herring is in fact one of the most abundant vertebrates on Earth, with schools comprising more than a billion individuals and an estimated global population in excess of 10^11^ fish (Feng *et al*., 2017). It is also one of very few marine species to successfully colonize the Baltic Sea, a brackish body of water formed after the last Ice Age, giving rise to the phenotypically distinct Baltic herring classified as a subspecies of the Atlantic herring.

Earlier work provided the first draft version of the herring genome (Martinez Barrio *et al*., 2016), and revealed regions with strong signals of selection related to both adaptation to the brackish Baltic Sea and differences in spawning time between herring populations (Martinez Barrio *et al*., 2016; Lamichhaney *et al*., 2017). In contrast, there is essentially no genetic differentiation at selectively neutral loci even between geographically distant populations, a fact documented already by isozyme and microsatellite analyses (Andersson *et al*., 1981; Larsson *et al*., 2010; Limborg *et al*., 2012; Ryman *et al*., 1984) and verified by whole genome sequencing and Fst analysis (Lamichhaney *et al*., 2012; Lamichhaney *et al*., 2017). However, while the signals of selection were strong, the fragmented nature of the draft genome made it challenging to determine the number of independent loci under selection as well as studying the impact of large-scale inversions and other structural variations.

Here, by combining a *de novo* long-read assembly of an Atlantic herring with long-range chromatin interaction information gathered via the Hi-C method (Lieberman-Aiden *et al*., 2009), we remedy this fragmentation and deliver a chromosome-level assembly of the herring genome comprising 26 autosomes with sizes ranging from 12.4 to 33.1 Mb and a total size of 726 Mb. We also show how this new assembly has a major impact on our ability to interpret the signals of selection. The final assembly version is publicly available via the European Nucleotide Archive (https://www.ebi.ac.uk/ena/data/view/GCA_900700415).

## RESULTS

### Compiling the hybrid assembly

The assembly is based on a new, *de novo*, assembly of an Atlantic herring, as opposed to a Baltic herring used for the previously published version 1.2 (Martinez Barrio *et al*., 2016). We generated 63 Gb (approximately 75x coverage) of sequence using Pacific Biosciences RSII cells and assembled the genome with FALCON-unzip (Chin *et al*., 2016). The FALCON-unzip assembly was processed through the PurgeHaplotigs pipeline (Roach *et al*., 2018), in order to remove redundant sequences from the primary assembly. This procedure resulted in a *de novo* assembly with a total size of 792.6 MB, a contig N50 of 1.61 Mb, comparable to the scaffold N50 (1.84 Mb) of the published v1.2 genome. Thus, the PacBio assembly achieves similar level of organization while eliminating a substantial degree of uncertainty, as v1.2 contains close to 10% undetermined bases (Ns) as compared to zero Ns in the FALCON-unzip assembly.

In order to obtain chromosome-level organization, a Hi-C library was prepared from liver and brain tissue from the same individual as used for the PacBio assembly, and it was sequenced on a BGISEQ-500 sequencer. Mapping with Juicer v1.5.6 (Durand *et al*., 2016b) yielded 99 million informative Hi-C-read-pairs, which were used to scaffold the PacBio *de novo* assembly into chromosome-level organization using the 3D-DNA workflow pipeline (Dudchenko *et al*., 2017) followed by manual correction using Juicebox v1.9.8 (Durand *et al*., 2016a). The output assembly was polished using Pilon v1.22 (Walker *et al*., 2014), based on 50x Ilumina paired-end coverage from the same individual. Finally, a custom R script was applied to eliminate a set of small, nearly identical repeats that were deemed likely to be redundant haplotypes based on analysis of the mapped read depth of a set of Ilumina short reads from a previously sequenced herring population (Martinez Barrio *et al*., 2016). This procedure eliminated in total 6.9 Mb (removed fragments are available as Supplementary Data S1). The entire assembly workflow is outlined in Figure 1. The resulting assembly version constitutes release v2.0.2, with the parts that could not be linked to the 26 chromosomes included as unplaced scaffolds. Summary statistics of the involved assemblies can be found in Table 1, while the size distribution of the assembled chromosomes is in Table 2. We assume that the 26 super-scaffolds correspond to chromosomes and have named these chr1-chr26 based on the size of the super-scaffold.

**Figure 1:**
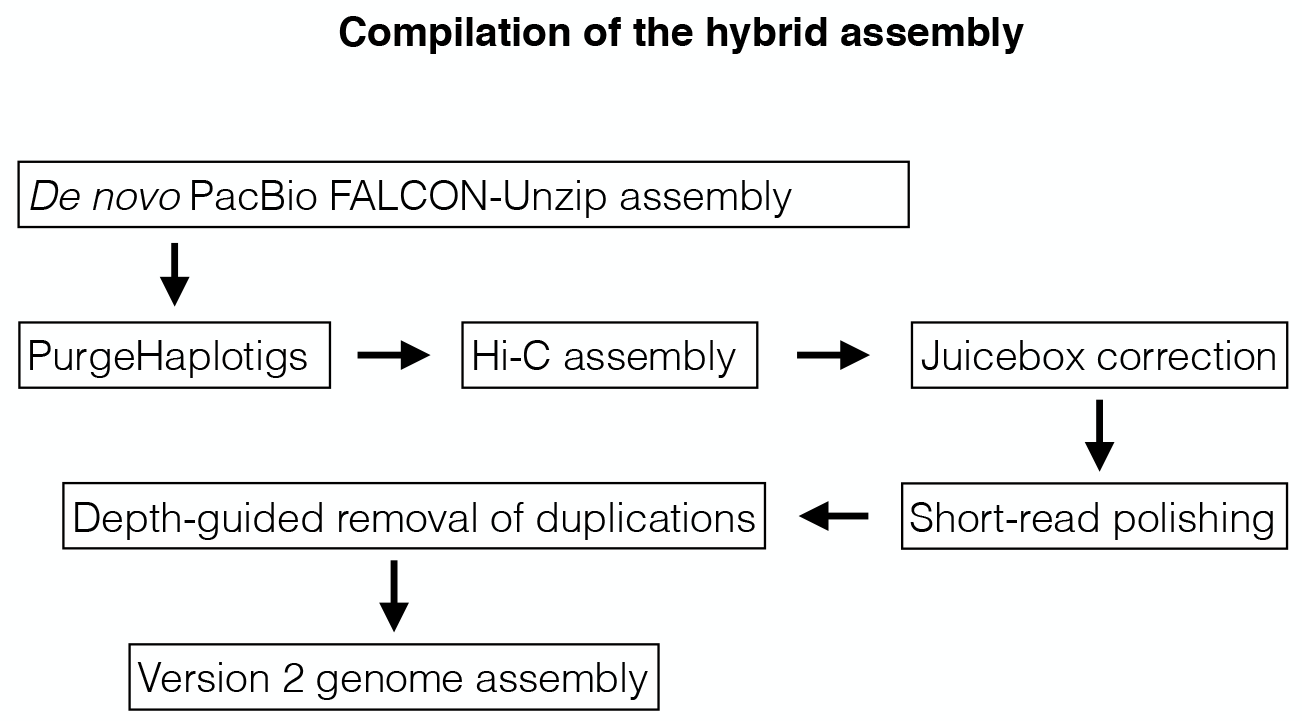
Flow-chart of the assembly compilation process.

**Table 1:** Summary statistics for different assemblies of the herring genome^1^.

|  | v1.2 | PacBio | v2.02 |
| --- | --- | --- | --- |
| Scaffolds | 6915 | - | 1723 |
| Scaffold N50 (Mb) | 1.84 | - | 29.85 |
| Scaffold L50 | 113 | - | 13 |
| Contigs | 73687 | 1356 | - |
| Contig N50 (kb) | 22.3 | 1615.0 | - |
| Contig L50 | 8566 | 119 | - |
| Size (Mb) | 808 | 793 | 726/786<sup>2</sup> |
| GC (%) | 44.1 | 44.2 | 44.1 |
| N (%) | 9.98 | 0.00 | 0.11 |

<sup>1</sup>v1.2 is the version published in Martinez-Barrio *et al.* (Martinez Barrio *et al.*, 2016), PacBio is the FALCON-unzip assembly presented here, v2.0.2 is the final hybrid assembly presented here.
<sup>2</sup>The 26 chromosomes total 726 Mb, unplaced scaffolds constitute 61 Mb.

**Table 2:** Physical and genetic sizes of the chromosomes.

| Name | Size (Mb) | Size (cM) |
| --- | --- | --- |
| Chr1 | 33.1 | 46.8 |
| Chr2 | 33.0 | 57.9 |
| Chr3 | 32.5 | 75.7 |
| Chr4 | 32.3 | 62.4 |
| Chr5 | 31.6 | 46.8 |
| Chr6 | 31.5 | 85.5 |
| Chr7 | 31.0 | 50.8 |
| Chr8 | 30.7 | 57.9 |
| Chr9 | 30.5 | 107.1 |
| Chr10 | 30.2 | 60.5 |
| Chr11 | 30.1 | 54.8 |
| Chr12 | 30.0 | 79.9 |
| Chr13 | 29.8 | 94.0 |
| Chr14 | 29.3 | 44.4 |
| Chr15 | 28.7 | 60.4 |
| Chr16 | 27.8 | 95.1 |
| Chr17 | 27.6 | 56.6 |
| Chr18 | 27.2 | 51.3 |
| Chr19 | 27.1 | 52.2 |
| Chr20 | 26.7 | 62.4 |
| Chr21 | 26.5 | 26.8 |
| Chr22 | 25.7 | 60.6 |
| Chr23 | 25.3 | 100.8 |
| Chr24 | 20.1 | 55.3 |
| Chr25 | 14.9 | 56.4 |
| Chr26 | 12.4 | 57.5 |
| Total: | 725.7 | 1659.9 |

### A comprehensive linkage map

We used pedigree data comprising two-full sib families, one Baltic herring and one Atlantic herring family, bred in captivity (Feng *et al*., 2017). The parents and the 45 Baltic and 50 Atlantic full-sibs were genotyped using a previously reported SNP chip (Martinez Barrio *et al*., 2016) comprising 41,575 markers across the genome. The markers formed 26 linkage groups in perfect agreement with the genome assembly and the linkage map confirmed the linear order of genomic segments along the chromosomes. The total length of the sex-average linkage map was 1,660 cM and the average recombination rate was 2.54 cM/Mb (Table 2, sex-specific maps in Supplementary Table S1); the male map was 13% longer than the female linkage map. The fact that the herring genome is composed of 26 chromosomes and the total map distance is 1,660 cM imply that there is slightly more than one recombination event per chromosome pair in each meiosis (26 x 50 cM = 1,300 cM). The chromosomal linkage maps showed a consistent pattern of non-uniform recombination rate across chromosomes, with the typical case being an L-shaped map where one section, in many cases 10 Mb or more in size, displays little to no recombination (Figure 2 *a*, *b*). In essence, it seems most chromosomes have one hot side and one cold side in terms of recombination rate. However, on the population level linkage disequilibrium (LD) still decays in the cold regions (data not shown), indicating that there is not a complete repression of recombination but rather a moderation of its frequency. The linkage map for chromosome 8 is shown in Figure 2*b*, and all maps are in Supplementary Figure S1.

**Figure 2:**
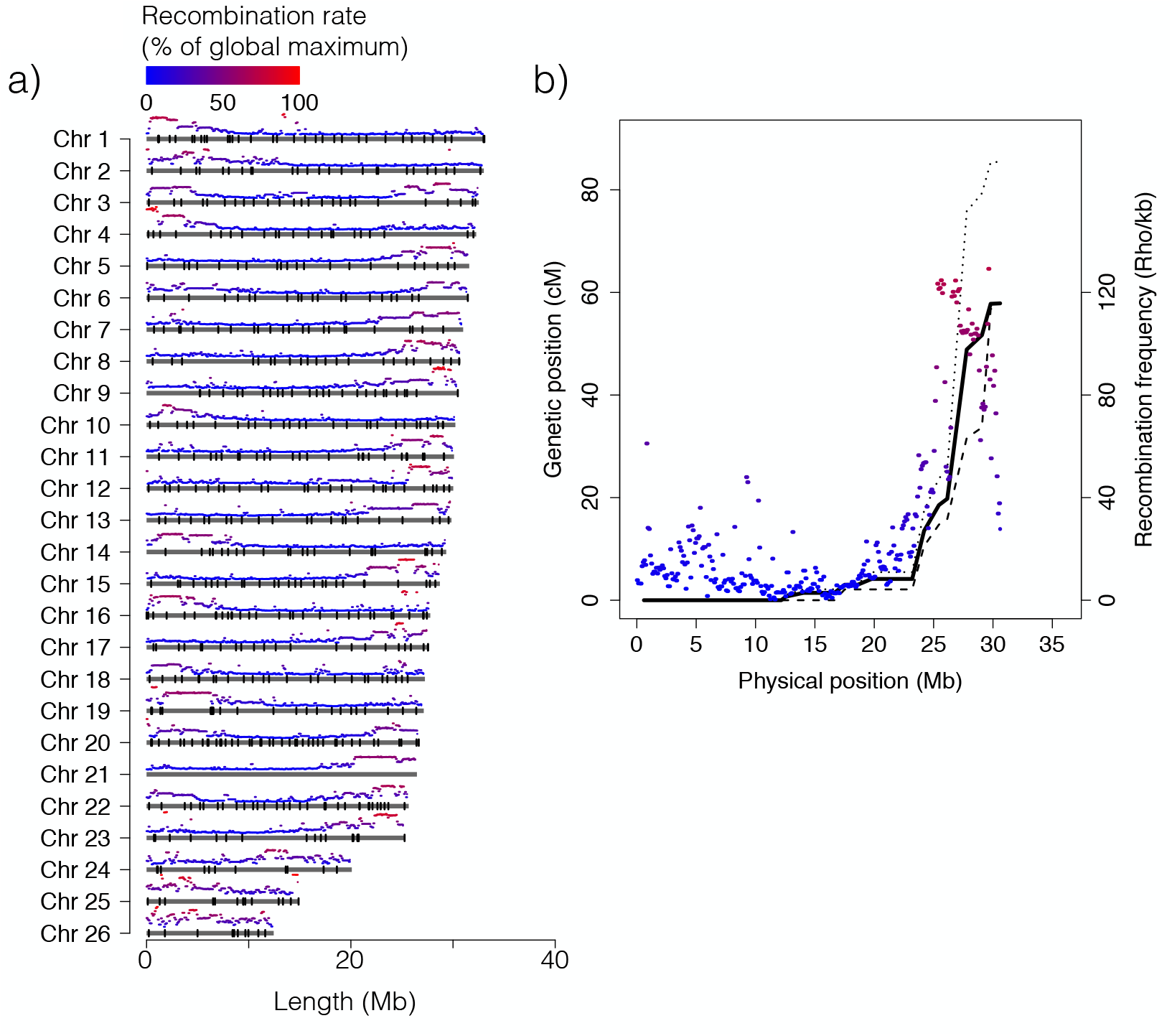
Chromosome size distribution and recombination rate. (*a)* Physical extent of the assembly for each chromosome, with average recombination rate, estimated using LD-hat (Auton & McVean, 2007), in 100 kb windows shown on top of each chromosome and markers used in the linkage map indicated as black bars. (*b)* Linkage map data (black lines) and recombination-rate profile (colored segments) estimated using LD-hat (Auton & McVean, 2007) for chromosome 8. Solid-line: sex-average linkage map, dashed line: male linkage map, dotted line: female linkage map.

### Chromosome-wise recombination profiles

The chromosomal-level assembly provides an opportunity to investigate the variation of recombination rates across the genome using population genetic data. Here, we constructed a recombination-rate profile in Atlantic herring based on patterns of linkage disequilibrium (LD) using LDhat (Auton & McVean, 2007). We estimated the crossover events among ∼1.7 million SNPs from 14 Baltic herring that have been individually sequenced (Martinez Barrio *et al*., 2016; Lamichhaney *et al*., 2017). A fine-scale recombination map was generated with mean recombination rate of ρ=31.3/kb, which corresponds to 2.1 cM/Mb, given a nucleotide diversity π = 0.3% (Martinez Barrio *et al*., 2016) and mutation rate μ = 2 × 10^-9^ determined by Feng *et al*. (Feng *et al*., 2017). Most chromosomes in this map display an “L” shape, in which one part of the chromosome has a higher recombination rate than the other (Figure 2*a*; Supplementary Figure S1). Thus, there was an excellent agreement between pedigree-based (linkage) and population-based (LD) estimates of the pattern of recombination along herring chromosomes (Figure 2*b*). Our estimation of the recombination rate in herring is close to the measure in zebrafish, which is about 1.6 cM/Mb (Bradley *et al*., 2011).

### Correspondence to karyotype and positioning of the centromeres

The version 2.0 genome assembly is consistent with the observed karyotype for the sister species Pacific herring (*Clupea pallasi*), both in terms of the number of chromosomes and their size distribution. Ida *et al*. (Ida *et al*., 1991) showed that the diploid genome consisted of 26 pairs (2*n* = 52), where most were of similar size with two smaller pairs that were speculated to be the results of a recent chromosome fission event. Out of the 26 pairs, three were determined to be metacentric or submetacentric, while the remaining 23 were reported to be acrocentric. Based on the recombination profile observed across the Atlantic herring chromosomes, we assume, based on the situation in many other species, that the recombination rate is relatively low towards centromeres and relatively high towards telomeres. Therefore, we suggest that chromosomes 3, 20 and 22 are metacentric, as the profile is shaped like a “U” rather than an “L” as is the case for most other chromosomes (Figure 2).

### Interchromosomal rearrangements are rare but intrachromosomal rearrangements are frequent among teleosts

The chromosome-level assembly allows comparisons of the herring genomic organization with that in other teleosts. We performed pair-wise whole genome alignments between the new herring assembly and eight other vertebrate species with chromosome-level assemblies publicly available: northern pike (*Esox lucius*), three-spine stickleback (*Gasterosteus aculeatus*), guppy (*Poecilia reticulata*), zebrafish (*Danio rerio*), medaka (*Oryzias latipes*), spotted gar (*Lepisosteus oculatus*), chicken (*Gallus gallus*), and humans (*Homo sapiens*). These comparisons revealed a very high degree of conserved synteny among teleosts, as illustrated by the comparison of the Atlantic herring and stickleback genomes (Figure 3*a*). However, the linear orders along chromosomes are highly rearranged (Figure 3*b*).

**Figure 3:**
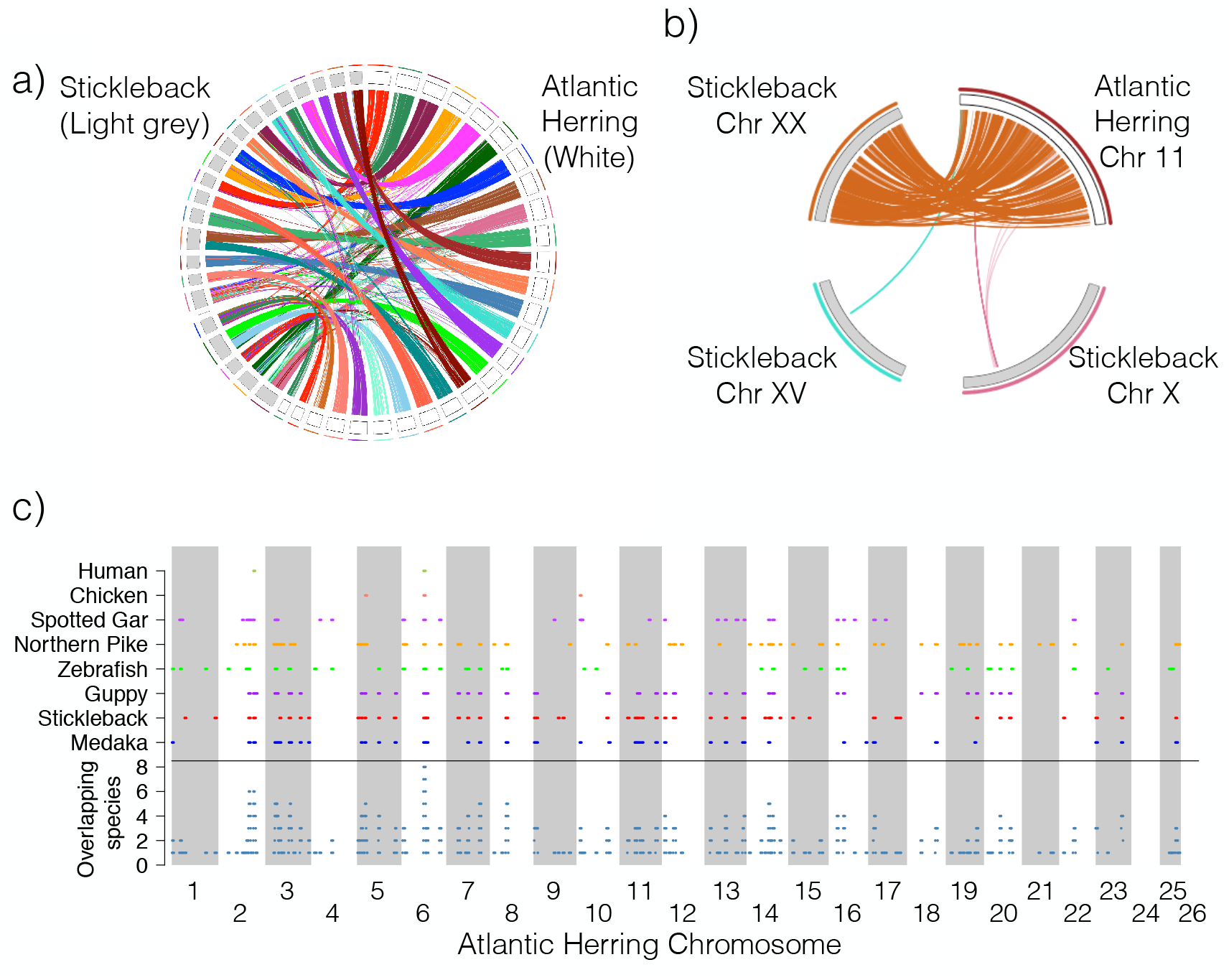
Conserved synteny among teleosts. (*a)* Whole genome alignment between the Atlantic herring and three-spine stickleback assemblies. (*b)* Detailed view of sequence homologies between herring chromosome 11 and stickleback chromosome XX, indicating many intra-chromosomal rearrangements. (*c)* Conserved co-linearity for a minimum of 1 Mb in the herring assembly across six teleost species, one bird (chicken) and one mammal (human). The section below the horizontal black line shows the number of species sharing a conserved region at all positions along the herring genome.

Additionally, we used SatsumaSynteny (Grabherr *et al*., 2010) to reveal collinear segments exceeding 1 Mb in size in pair-wise comparisons among the nine species of vertebrates including the Atlantic herring (Figure 3*c*). By intersecting the positions of such co-linear fragments, we note that the amount of overlap by far exceeds expectation, under a model that the distribution of these regions is independent among the different species pairs, for all numbers of overlapping species greater than three (Supplementary Figure S2*a*). This expectation is based on the extent of the co-linear regions of 1 Mb or larger in each pairwise comparison, which ranges from ten to five percent when comparing Atlantic herring to other teleosts to less than one percent in comparisons with chicken and human. If the outcome of all comparisons were independent, we do not expect any regions where all nine species have a co-linear block, whereas the observed case is one such block, spanning 1.66 Mb on chromosome 6 in herring co-ordinates and corresponding to a 5.5 Mb region on human chromosome 16 (Supplementary Figure S2*b*). For both the chicken and humans there are only two co-linear, as compared to the herring assembly, blocks detected across the genome, and in both cases one of the two is located at this region. In the teleosts, there is an additional region on chromosome 2 that is shared, as well as a few that are shared in six out of seven species. In addition to the amount of overlapping sequence, the endpoints of the overlapping blocks are often closely aligned, which is not expected under a model where re-arrangements occur at arbitrary locations. Finally, the pattern is not driven by rearrangements unique to the herring genome or caused by errors in the current assembly. An analogous analysis among the other species involved show there are many rearrangements between each pair included in the study (data not shown). GO analysis of the genes inside the co-linear blocks conserved in six or more species did not reveal any striking enrichment, leading us to conclude that the likely reason for non-random re-arrangements, and consequential maintenance of these blocks intact across large time-spans, lies in long-distance regulatory elements rather than grouping of functionally related genes.

### The number of independent signals of selection associated with ecological adaptation

Our previous data indicated that adaptation to the brackish Baltic Sea, and to different spawning times (spring-autumn) were both influenced by a large number of independent loci (Martinez Barrio *et al*., 2016). In the scaffold-level assembly, both contrasts yielded hundreds of separate signals. This number was known to be an overestimate, due to the fragmented nature of the previous assembly, but the degree of overestimation was unclear. Now, by transferring the same dataset to the new assembly and defining an independent peak as (i.) at least two SNPs with χ^2^-test *p* values less than 10^-20^, the same threshold as used in Martinez-Barrio *et al*. (Martinez Barrio *et al*., 2016), (ii.) spanning at least 100 bases and (iii.) separated from the next peak by at least 100 kb, we find 125 independent loci associated with adaptation to the Baltic Sea, and 22 loci associated with different spawning times (Figure 4 *a*,*b*). Using a lower cut-off of 10^-15^, yields 195 and 47 independent loci, respectively, indicating that, in qualitative terms, the result is not sensitive to the choice of cut-off value. It is clear that while both these adaptations have left a complex genomic footprint, there is a 4- to 5-fold difference in the number of loci reaching statistical significance in our data. The results make sense because adaptation to the Baltic Sea appears to be a more complex process than adaptation to different spawning times; the Baltic Sea differs from the Atlantic Ocean as regards salinity, optic characteristics, depth, higher seasonal variation in temperature, plankton production, predators present and pollution.

**Figure 4:**
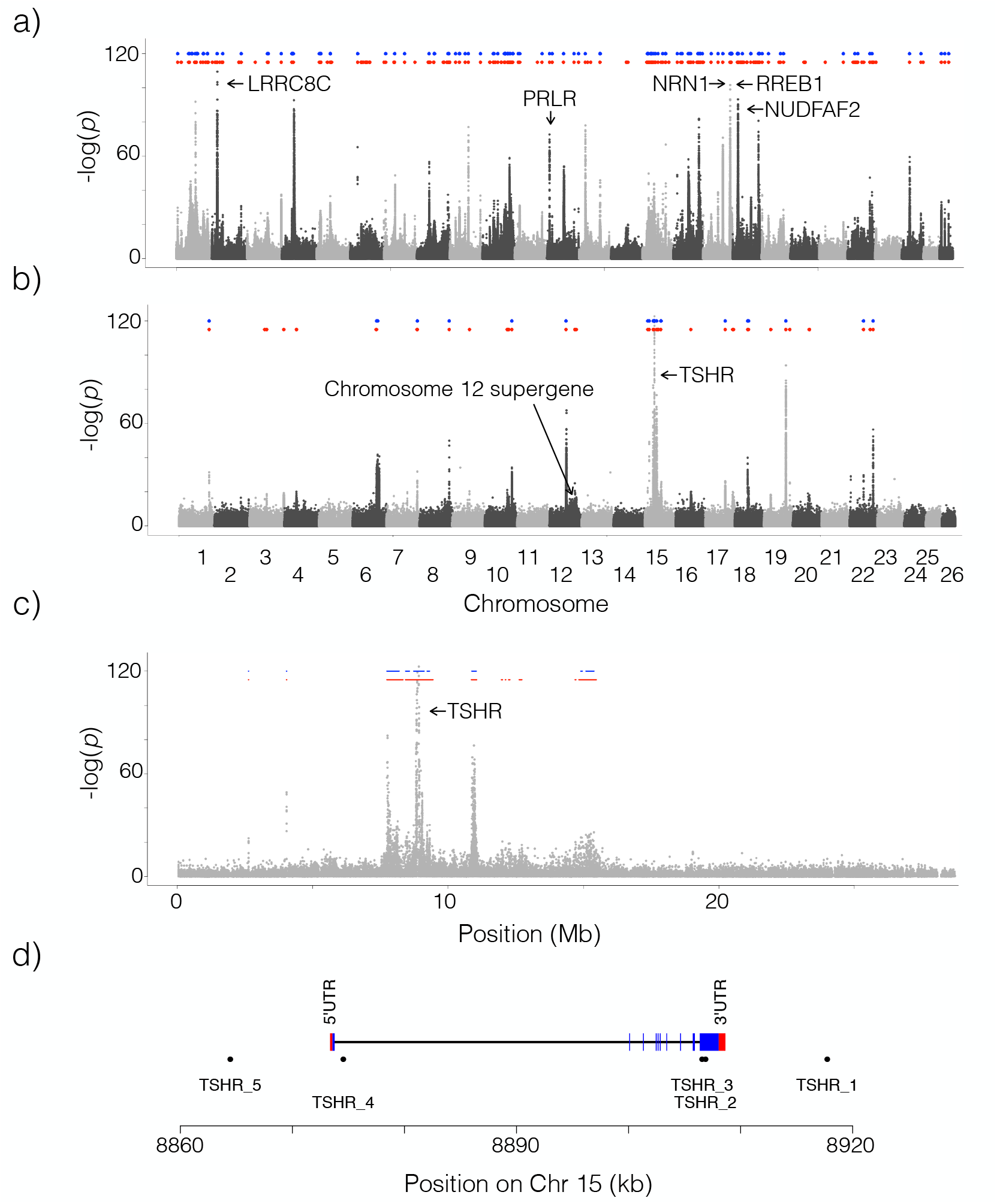
Signals of selections associated with ecological adaptation. The panels show the association, measured as –log(*p*) from χ^2^-tests on read-counts of previously published data (Martinez Barrio et al., 2016) re-plotted along the new assembly. (*a)* Genetic differentiation between seven populations of Atlantic herring from the Atlantic Ocean, North Sea, Skagerrak and Kattegat (salinity in the range 20-35 psu) and ten populations from the Baltic Sea (salinity in the range 3-12 psu). (*b)* Genetic differentiation between ten populations of spring-spawning herring versus three populations of autumn-spawning herring. In both panels, the blue and red dots indicate identified, independent regions of selection at a p-value cut-off of either 10^-20^ (blue) or 10^-15^ (red). (*c)* Zoom in on chromosome 15 for the contrast based on differences in spawning time, which contains the most significant peak, located around *TSHR*. (*d)* Improved gene model of *TSHR*. TSHR_1 to 5 are selected SNPs showing highly significant association (P < 10^-95^) to spawning time and/or being non-synonymous coding, while covering the extent of the *THSR* gene model.

### Resolving spawning time-associated variation at the *TSHR* locus

Our previous work (Lamichhaney *et al*., 2017; Martinez Barrio *et al*., 2016) showed that the SNPs most strongly associated with variation in spawning time are located immediately adjacent to the gene coding for the thyroid-stimulating hormone receptor (*TSHR*) (Figure 4*c*). However, the gene model in the annotation of the v1.2 genome was truncated compared to homologues from other species. In order to improve the gene model to properly interpret the effects of the associated SNPs, we performed 5’ and 3’ RACE experiments, and can now present a complete annotation of the herring *TSHR* gene (Figure 4*d*). This shows that the five SNPs denoted TSHR_1 to −5, which span the strongest genome-wide association peak to spawning time, are located within TSHR or within 15 kb upstream or downstream of the gene (Figure 4*d*).

### Identification of a supergene on chromosome 12

The pattern of linkage disequilibrium (LD) across the herring genome were analyzed in more than 1,170 individuals collected as part of a high school project (Forskarhjälpen) where the students collected samples around the Swedish coast. The samples were genotyped for approximately 45,000 SNPs. This analysis revealed an extensive LD block spanning approximately 7.8 Mb on chromosome 12, from 17.8 Mb to 25.6 Mb, that in the old assembly was divided into four different scaffolds. This region also stood out in different screens for ecological adaptation since the genetic differentiation was more or less equally strong across the block (Figure 5*a*), lacking the typical pyramidal peak shape. Inside the block there was no correlation between the strength of LD and physical distance (Figure 5*b*, top). This is in contrast to the genomic background, where the LD decay is very sharp – typically *r*^2^-values decay to 0.1 or less within a few kb. This finding indicated the presence of a possible supergene, either in the form of an inversion or a block of otherwise repressed recombination. However, the pattern contained inconsistencies, specifically there were moderately linked markers interspersed with virtually perfectly linked ones across the full extent of the block.

**Figure 5:**
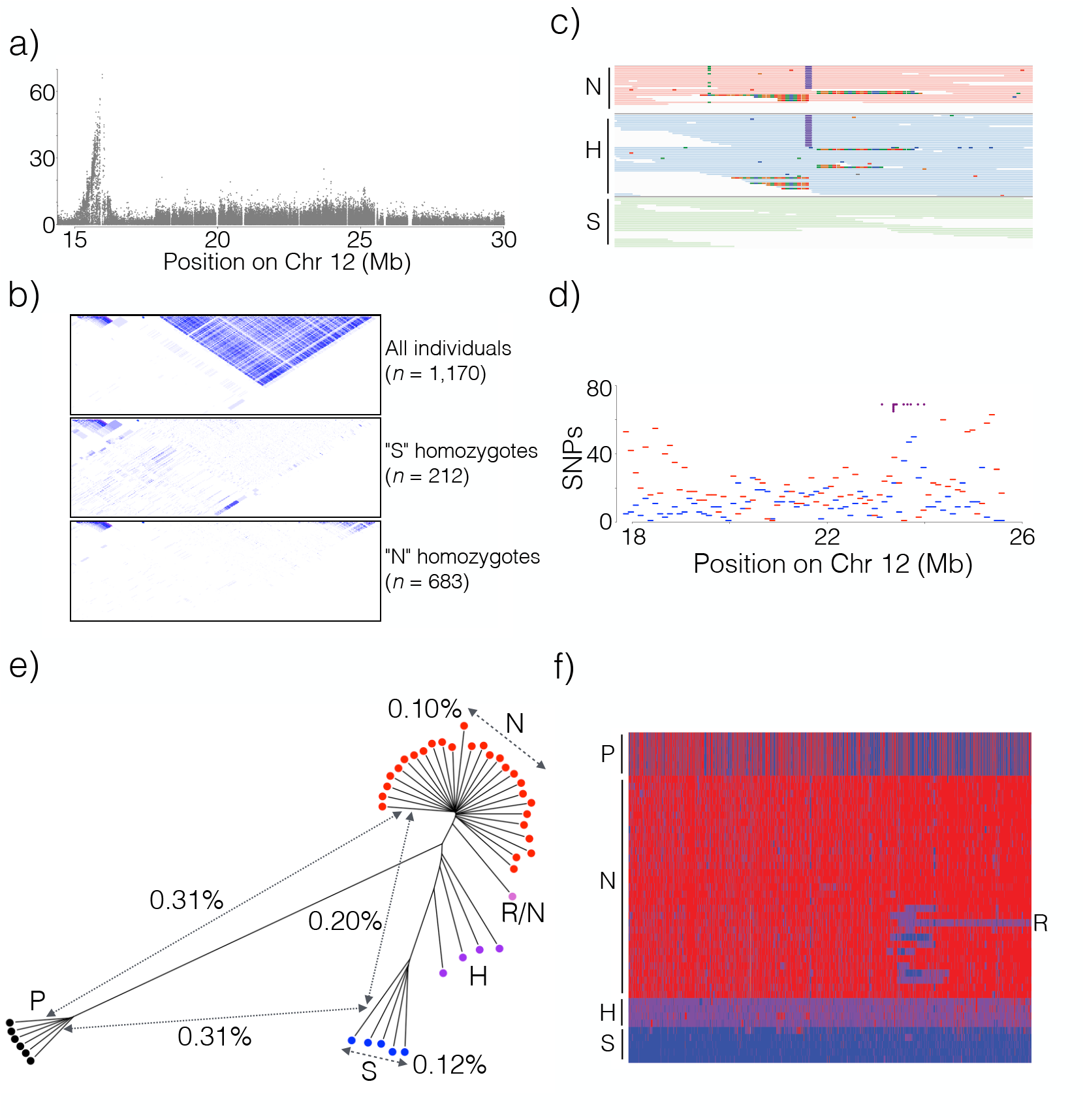
Identification of a 7.8 Mb inversion on herring chromosome 12. (*a)* The spawning time contrast for Chromosome 12 highlight the block-like association pattern for the region from 17.9 to 25.6 Mb. (*b)* LD patterns across the region in different groups of individuals sorted according to genotype for the putative inversion. (*c)* Read alignments across the inversion breakpoint at Chr12:17.9 Mb. The letters designate the supergene-type of the individual as follows: N = Northern homozygote; S=Southern homozygote; H = N/S heterozygote. (*d)* The distribution, as number of SNPs per 100 kb, of shared (blue) and diagnostic (red) SNPs across the inversion region. Purple dots (top) are estimated locations of breakpoints in individuals that appear to carry a recombinant chromosome (see text). (*e)* Neighbor-joining tree based on genotypes for all SNPs in the inversion region called from individual whole genome sequencing. The distances indicated across the tree are average nucleotide differences between haplotypes, either within groups (dashed) or between groups (solid). Letters are used as in (*c*), with the following extensions: R/N = individual carrying an N haplotype and a recombinant haplotype (see text); P=Pacific herring. (*f)* Heatmap of the genotypes for diagnostic SNPs based on individual whole genome sequencing. Supergene type of the samples is indicated as in (*e*).

Led by the LD patterns revealed in the population data, detailed examination of the pedigree data used to construct the linkage map revealed an elevated frequency of heterozygous SNPs in one out of four parents across the putative inversion, a pattern that was repeated in 28 out of 50 of the offspring in that family, consistent with proper Mendelian inheritance. Assuming that the high-heterozygosity parent and offspring were, in fact, heterozygous for the proposed inversion allowed us to deduce supergene haplotypes and use these to genotype all individuals in the pedigree. This, in turn, made it possible to genotype even unrelated individuals, based on their similarity, measured as an average of the SNP genotypes over the region, to the two reference haplotypes; these were denoted Northern (N) and Southern (S) based on their geographic distribution (see below). We applied this to the set of 1,170 fish collected from 20 sites around the coast of Sweden. Finally, by analyzing LD patterns in the groups of individuals determined to be homozygous for different haplotypes, we noted a lack of strong LD across the entire region in both groups (Figure 5*b*, middle and bottom). This indicates that recombination occurs between chromosomes of the same class but is strongly repressed in heterozygotes, which is the expected pattern for an inversion but not if recombination is generally suppressed in the region for other reasons.

We carefully examined reads from whole genome sequence data from individuals with different genotypes in an attempt to identify inversion break points. We identified inverted repeat patterns at both ends of the putative inversion block (Supplementary Figure S3). For the putative breakpoint at 17.9 Mb, we found short-read mismatch patterns at the edge of this repeat (position 17,826,318 bp) that correlated perfectly with the SNP-based supergene genotype (Figure 5*c*). No short-read mismatches (e.g. soft-clipped reads) was identified in individuals classified SS homozygotes, consistent with the fact that the reference assembly is based on the S haplotype. In contrast, 50% and 100% of the short reads from NS heterozygotes and NN homozygotes, respectively, were mismatched at this position. Thus, we consider this as a putative inversion breakpoint, but at present we cannot exclude the possibility that this is a structural variant in complete LD with the true breakpoint. We were not able to identify similar mismatched reads at the other breakpoint (around 25.6 Mb) due to the high repeat content. Taken together, these observations support the hypothesis that the extremely strong LD in this region is in fact caused by an inversion and that the inverted repeats have facilitated its occurrence.

Examining individual SNP allele frequencies in each group of locus-wide homozygotes, we were able to determine that a fraction of the SNPs within the interval were shared between the two haplotype-groups. This is not expected of a canonical inversion with a single origination event and with complete suppression of recombination. Thus, some amount of genetic exchange must be ongoing between the two haplotypes. We identified diagnostic SNPs that were essentially fixed for different alleles in the two haplotype-groups and shared SNPs that were segregating in both groups. As indicated in Figure 5*d*, the relative proportion of diagnostic (red) and shared (blue) SNPs varies along the region with an enrichment for diagnostic SNPs at each end of the interval, and a peak of shared SNPs, and corresponding lack of diagnostic SNPs, around position 23.5 Mb. Supplementary Figure S4 shows the allele frequency distributions among the 11,965 typed SNPs in “S” and “N” homozygotes. The higher number of SNPs close to fixation (MAF < 5%) in the “N” group indicates that it is the derived version, which could be correlated to northward expansion in response to receding glaciation.

This analysis also revealed a rare class of individuals (12 out of 1170) that appear to carry a third haplotype where the segment 17.8-23.4 Mb follows the “Southern” haplotype, while the block from 23.4-25.6 Mb is consistent with the “Northern” haplotype. The estimated switching-points of these 12 individuals are shown as purple dots in Figure 5*d*. There is also a residual long-range LD among “Northern” homozygotes over the same sub-region (Figure 5*b*). Interestingly, the end of this region is close to the region where the peak for the proportion of shared SNPs is located.

We constructed a genetic distance tree for the supergene region based on individual whole genome sequence data (Figure 5*e*); the color of each leaf in the tree corresponds to the supergene-type. This shows the expected clustering of the two supergene-types (N or S), while the heterozygotes (H) are positioned in between. Notably there is one individual that carries one copy of the partial inversion haplotype (R) discussed above, with an estimated breakpoint in exactly the same region as predicted for the 12 out of 1,170 individuals genotyped using the SNP chip. The tree also reveals that the inversion must have occurred well after the divergence from the Pacific herring, because the two alleles are equidistant from alleles found in Pacific herring (Figure 5*e*). The nucleotide diversity inside each haplotype group is lower than the genomic average of 0.3%, which is consistent with both lower effective population size, due to restricted recombination, for this region compared to the rest of the genome, as well as a bottleneck when the inversion was formed.

A heat map based on diagnostic SNPs, deduced from individual whole genome sequence data, from the inversion region support the notion that the two haplotype groups evolved subsequent to the split between Atlantic and Pacific herring and illustrates the extreme LD across the region (Figure 5*f*). Interestingly, this analysis provides strong evidence for ongoing recombination between haplotypes in the interval from 23.1 to 24.0 Mb. Additionally, the heatmap makes it clear that the individual labeled “R/N” in Figure 5*e* carries one copy of the recombinant haplotype found in 12 of the SNP-typed individuals, with partial Northern (17.8-23.4 Mb) and Southern (23.4-25.6 Mb) configuration.

### The supergene on chromosome 12 is underlying ecological adaptation

Using the SNPs found to be essentially fixed for different alleles in the Northern and Southern haplotype groups, *i.e.* those SNPs found in the lower-right corner of Figure 6*a*, we traced the frequency of each haplotype also in pooled samples, by using the average allele frequency at these diagnostic SNPs as a proxy for the frequency of these haplotypes. The frequencies obtained with this method were consistent with the individual classification.

**Figure 6:**
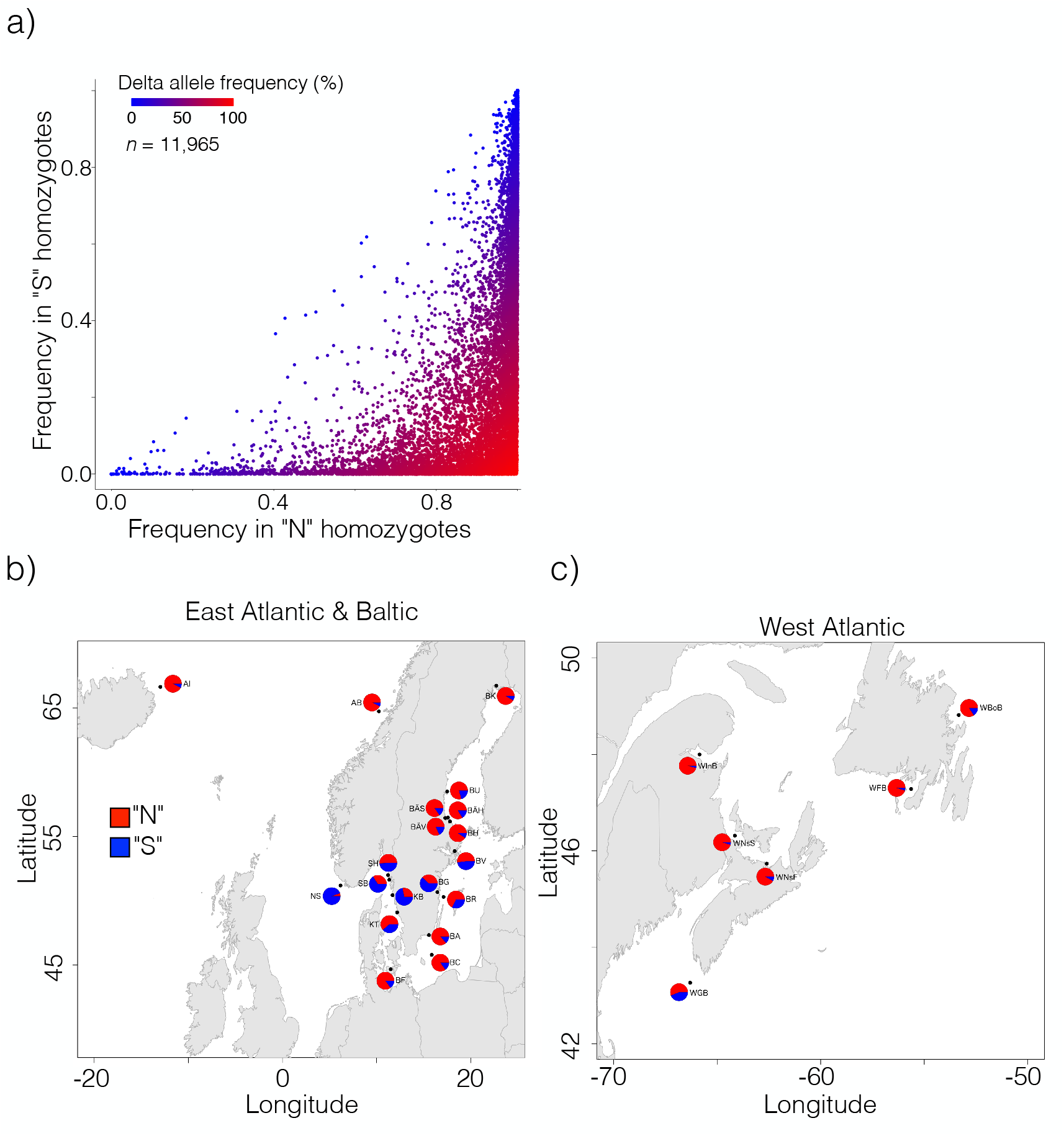
Genetic differentiation in the region encompassing the putative inversion on chromosome 12. (*a)* Allele frequencies for all SNPs (*n*=11,965) inside the inversion among individuals homozygous for the Southern (S) (y-axis) or Northern (N) haplotypes (x-axis). All frequencies are expressed in terms of the allele that is more common in the “N-”context than in the “S-”context. (*b*, *c)* Estimated allele frequencies for the Northern and Southern haplotypes in pooled population samples in the Baltic Sea and East Atlantic (*b*) and in the West Atlantic (*c*). The location and date of capture of the pooled samples are listed in Lamichhaney *et al*. (Lamichhaney *et al*., 2017).

The 7.8 MB region contains 225 genes, with an additional approximately 10 genes located in flanking positions on the outside of the estimated breakpoint positions. Given the size of the region, it is difficult to determine which genes and or variants that contribute to the fitness effects of the inversion, but, based on the disruption of the local context, genes near the breakpoints are more likely to have their functionality affected. Thus, the genes closest to the two breakpoints are listed in Supplementary Table S2.

The estimated haplotype frequencies in the pooled data, which covers a wide range of herring populations, revealed a highly significant genetic differentiation among populations (Figure 6*b*,*c*). There was a consistent trend in West Atlantic, East Atlantic and in the Baltic Sea that the populations spawning most northerly had a very high frequency of the Northern haplotype while the Southern haplotype dominated in populations spawning more southerly. The only exception to this trend was a few populations in Southern Baltic Sea that had a high frequency of the Northern haplotype. The most extreme population almost fixed for the Southern haplotype was the one representing autumn-spawning North Sea herring (NS in Figure 6*b*). This strong genetic differentiation is never observed at selectively neutral loci among the populations included in this analysis (Lamichhaney *et al*., 2017; Martinez Barrio *et al*., 2016), which provide conclusive evidence that this supergene polymorphism must be under selection, possibly related to temperature at spawning, which is known to be a major stressor for the southernmost and high temperature exposed herring populations (Ojaveer *et al*., 2015; Peck *et al*., 2012).

## DISCUSSION

### Integrity of the assembly

The overall organization of the assembly presented here is mainly supported by two separate observations: the one-to-one correlation between putative chromosomes and independently determined linkage groups, and the discrete blocks detected in the Hi-C contact map. Based on these two datasets, we are confident that the 26 super-scaffolds present in the assembly match the 26 physical chromosomes identified by karyotyping of the Pacific herring (Ida *et al*., 1991), a sister taxa to the Atlantic herring. Furthermore, the high quality of the assembly is strongly supported by the very high degree of conserved synteny between our Atlantic herring genome and those of other teleosts (see below).

### Inter-chromosomal rearrangements are rare while intra-chromosomal rearrangements are abundant in teleosts

This study revealed a remarkable contrast between conserved synteny with very few interchromosomal rearrangements between Atlantic herring and even distantly related teleosts, and the frequent occurrence of intrachromosomal rearrangements. This finding is consistent with the results of previous comparisons among teleost genomes (Amores *et al*., 2014; Rondeau *et al*., 2014). There is an interesting difference among vertebrate groups where fishes and birds usually show few interchromosomal rearrangements but frequent intrachromosomal rearrangements. In contrast, there is an opposite trend among mammals where often fairly closely related species like mouse and rat show many interchromosomal rearrangements (Coghlan *et al*., 2005). The explanation for this difference between vertebrate groups is still unknown. It has been proposed that differences in repeat contents could play a role (Amores *et al*., 2014), since repeats may facilitate non-orthologous recombination. However, this should affect both inter- and intra-chromosomal rearrangements and does not appear to explain this difference in the relative frequency of the two types of rearrangements among different vertebrates.

The probability that a chromosomal rearrangement becomes fixed in a species will depend on how the rearrangement affects fitness, and both positive and negative selection may occur. It can do this in two ways, one is that the rearrangement may affect gene regulation by changing chromatin organization. This is, for instance, well illustrated by mutations affecting comb morphology in chickens that are all caused by structural rearrangements leading to ectopic expression of transcription factor genes (Dorshorst *et al*., 2011; Imsland *et al*., 2012; Wright *et al*., 2009). The other way is that in heterozygous carriers, the rearrangement *per se* may increase the occurrence of unbalanced gametes and therefore cause reduced fertility. Translocation heterozygotes in particular often show impaired fertility while small inversions (<30% of a chromosome arm) has small or no negative effect on fertility (Morin *et al*., 2017). Thus, an explanation for the difference in the relative abundance of inter- and intra-chromosomal rearrangements among vertebrate groups could be due to differences how the two types of rearrangements affect meioses. The probability that a chromosomal rearrangement, despite a negative effect on fertility, goes to fixation also depends on population structure because it is more likely to happen due to genetic drift in a small population than in large panmictic populations like those in the Atlantic herring. The very high degree of conserved synteny among teleosts shows that interchromosomal rearrangements have not been an important mechanism promoting speciation and diversification in fish.

Despite the common occurrence of intrachromosomal rearrangements among teleosts we noted a highly non-random pattern where chromosomal breakpoints occur, leaving megabase regions where the linear order is unbroken across species, sometimes also including conservation of gene order in avian and mammalian genomes. This is most likely caused by the presence of long-range regulatory domains and co-regulated clusters of genes.

### Large-scale inversions and their evolutionary significance

In our previous studies (Lamichhaney *et al*., 2017; Martinez Barrio *et al*., 2016), using the scaffold-level assembly available at the time, it was indicated that small inversions were not a major contributor to differences between herring populations while the issue of larger, megabase-scale, inversions was intractable given the scaffold length distribution. Using the improved assembly presented here, we now have clear evidence of selection acting on a supergene that is, in fact, a 7.8 Mb inversion. We were even able to identify a putative inversion breakpoint at position 17,826,318 bp on chromosome 12. However, our data also illustrates how difficult it is to exactly define inversion breakpoints because they are often embedded in repeat regions.

Our observations fit with a supergene-model, where large essentially non-recombining haplotypes allow accumulation of multiple causal variants synergistically affecting fitness. The divergence time between the two haplotype groups (Figure 5*e*) cannot be accurately determined due to the ongoing genetic exchange between haplotype groups. However, the fact that the observed nucleotide divergence (0.2%) is lower than the genomic average (about 0.3%) indicates that the putative inversion may be fairly recent. The fact that the Pacific herring carries a separate haplotype limits the maximum age of the inversion to after the split of the two species. An interesting finding was that there is apparently ongoing genetic exchange between the inversion alleles as illustrated by the presence of many shared polymorphisms as well as recombinant haplotypes (Figure 5*f*). It is likely that both double recombination and gene conversion contributes to this genetic exchange. It is in fact a characteristic feature of inversions that some recombination occurs between alleles although recombination is severely suppressed. This has for instance been well documented for the inversion underlying the Rosecomb phenotype in domestic chicken (Imsland *et al*., 2012) and the inversion associated with variant mating strategies in the ruff (Kupper *et al*., 2016; Lamichhaney *et al*., 2016).

While it is currently not known what is the cause of fitness differences between the inversion variants in herring, it appears highly likely that it is related to ecological adaptation in relation to the water temperature during gonadal maturation before spawning or the water temperature at spawning/early larval development. There is a clear clinal variation of an increasing frequency of the Southern variant from north to south both in East and West Atlantic (Figure 6).

The supergene on herring chromosome 12 underlying ecological adaptation adds to the growing list of supergenes associated with morphological variation and ecological adaptation (Schwander *et al*., 2014). The first very early examples of supergenes under balancing selection were those detected by cytogenetic studies of polytene chromosomes in *Drosophila* (Dobzhansky & Sturtevant, 1938). More recent examples include supergenes controlling mimicry in butterflies (Zhang *et al*., 2017), social behavior in fire ants (Wang *et al*., 2013), plumage variation and mating preferences in white-throated sparrows (Tuttle *et al*., 2016) and alternative male mating strategies in ruff (Kupper *et al*., 2016; Lamichhaney *et al*., 2016). Furthermore, five putative inversions ranging in size from 3.5 to 18.5 Mb have recently been associated with migratory behavior and geographical distribution in the Atlantic cod (*Gadus morhua*) (Berg *et al*., 2017). The present study shows how chromosome-based assemblies will facilitate the identification of many other similar examples of supergenes.

## METHODS

### FALCON *de novo* assembly

Genomic DNA was fragmented to 20 kb using a DNA shearing device (Hydroshear, Digilab), and the sheared fragments were size-selected for the 7-50 kb size range using Blue Pippin (Sage Science). The sequencing library was prepared following the standard SMRT bell construction protocol (PacBio). The library was sequenced on 100 PacBio RSII SMRT cells using the P6-C4 chemistry. Raw data was imported into SMRT Analysis software 2.3.0 (PacBio). Subreads shorter than 500 bp or with a quality (QV) <80 were filtered out. The final data set contained 63.1 Gb of filtered subreads with N50 of 15.6 kb and was used for *de novo* assembly with Falcon (pb-falcon 0.2.4) (Chin *et al*., 2016). To further improve the assembly we ran Falcon Unzip (pb-falcon 0.2.4) (Chin *et al*., 2016) followed by consensus calling using the Arrow algorithm. In order to remove highly heterozygous haplotypes assembled as separate primary contigs, we ran the Purge Haplotigs pipeline (v1.0.4) (Roach *et al*., 2018), which identifies and reassigns allelic contigs. The configuration file used for assembly constitutes Supplementary Text S1.

### Hi-C-library construction and sequencing

*In situ* Hi-C was conducted following the protocol provided by Rao *et al*. (Rao *et al*., 2014) with minor modifications. The restriction endonuclease MboI was used to digest DNA, followed by biotinylated residue labeling. The Hi-C library was then sequenced on BGISEQ-500 platform with pair-end sequencing using a read length of 50 bp. The raw number of reads was 656,695,125, out of which 98,838,909 provided useful H-iC contact information.

### Annotation

The herring gene set was generated via the Ensembl Gene Annotation (Aken *et al*., 2016). Annotation was created primarily though alignment of short read RNA-seq data to the genome, with gap filling via protein- to-genome alignments of a select set of vertebrate proteins from UniProt (UniProt, 2019). The short-read RNA-seq data were sourced from samples generated as part of this project (PRJEB31270) along with publicly available data from a Baltic Herring individual fished at Timraroe (PRJNA179110). The UniProt vertebrate proteins used had experimental evidence for existence at the protein or transcript level.

At each locus, low quality transcript models were removed and the data were collapsed and consolidated into a final gene model plus its associated non-redundant transcript set. When collapsing the data, priority was given to models derived from transcriptomic data. For each putative transcript, the coverage of the longest open reading frame was assessed in relation to known vertebrate proteins, to help differentiate between true isoforms and fragments. In loci where the RNA-seq data were fragmented or missing, homology data took precedence, with preference given to longer transcripts that had strong intron support from the short-read data.

Gene models from the above process were classified into three main types: protein-coding, pseudogene, and long non-coding. Models with hits to know proteins, and few structural abnormalities (i.e. they had canonical splice sites, introns passing a minimum size threshold, low level of repeat coverage) were classified as protein-coding. Models with hits to known protein, but having multiple issues in their underlying structure, were classified as pseudogenes. Single-exon models with a corresponding multi-exon copy elsewhere in the genome were classified as processed pseudogenes. If a model failed to meet the criteria of any of the previously described categories, did not overlap a protein-coding gene, and had been constructed from transcriptomic data then it was considered as a potential lncRNA. Potential lncRNAs were filtered to remove transcripts that did not have at least two valid splice sites or cover 1000bp (to remove transcriptional noise).

A separate pipeline was run to annotation small non-coding genes. Putative miRNAs were identified via a BLAST (Altschul *et al*., 1990) of miRBase (Kozomara *et al*., 2019) against the genome, before passing the results in to RNAfold (Gruber *et al*., 2008). Poor quality and repeat-ridden alignments were discarded. Other types of small non-coding genes were annotated by scanning Rfam (Kalvari *et al*., 2018) against the genome and passing the results into Infernal (Nawrocki & Eddy, 2013).

The annotation for the Atlantic herring annotation will be made available as part of Ensembl release 98 (expected October 2019). A detailed description of the annotation pipeline is available as Supplementary Text S2

### SNP chip genotyping

We previously designed a 60k Affymetrix SNP chip (Martinez Barrio *et al*., 2016). For this study, this SNP chip has been used to genotype two data sets: (i) A pedigree comprising two families with two parents and approximately 50 offspring each. (ii) 1,170 individuals collected in the school project “Forskarhjälpen” in which students from 20 junior high schools from across Sweden contributed to research by collecting a sample of approximately 50 herring from one locality per school (Supplementary Table S3).

### Construction of a high-resolution linkage map

The linkage map was constructed using a pedigree material comprising two large full-sib families, one generated using a male and a female Baltic herring from Sweden and the other using a male and a female Atlantic herring from Norway (Feng *et al*., 2017). Fifty full-sib progeny from the Atlantic family and 45 from the Baltic family were genotyped for about 45,000 SNPs using our previously described SNP-chip (Martinez Barrio *et al*., 2016). Linkage groups spanning the herring genome were constructed using these data and the LepMap 2 (Rastas *et al*., 2013) software. The raw linkage groups were in overall concordance with the Hi-C-assemblies, allowing each chromosome to be conclusively associated with a distinct linkage group. Based on this association, we were able to prune the marker set based on physical position, with the intent of controlling artificial map expansion while marinating coverage across the entire chromosome. The final, ordered, linkage maps of each chromosome were calculated using CriMap2.5 (Green *et al*., 1990), and the locations of the markers used are shown in Figure 2 with detailed versions found in Supplementary Figure S1 and sex-specific maps found in Supplementary Table S1.

### Linkage disequilibrium analysis (LD) and chromosome-wise recombination profiles

LD between markers was measured as correlation between genotypes, calculated using the “r2fast” method from the R-package GenABEL (Aulchenko *et al*., 2007). To estimate the recombination rates from population data, we applied the LDhat v2.2 package (Auton & McVean, 2007) on genetic markers from 14 Baltic individuals phased using Beagle4.0 (Browning & Browning, 2007). The expected crossover events (*ρ*) between each pair of neighboring markers were calculated with the interval program for 1,000,000 iterations of rjMCMC procedure with sampling every 2,000 iterations. The first 50 iterations were discarded as burn-in. The block penalty 5 was determined after comparing output from simulations with block penalties of 5, 20, 50 and 100. The population recombination map was summarized by summing up the *ρ* from every 100 kb window, and only windows containing 50 to 2,000 variable sites were included in the final map.

### Improvement of the TSHR gene model

Total RNA was extracted from the eye of a spring-spawning Atlantic herring using the RNeasy Mini Kit (Qiagen). Six μg of the isolated RNA was used for 5’- and 3’ RACE reactions with a FirstChoice^TM^ RLM-RACE Kit (ThermoFisher Scientific) according to the manufacturer’s protocol. Nested RACE PCRs were performed in a 25 μL reaction containing 5X KAPA2G Buffer B, 0.24 mM dNTPs, 0.5 μM each of the forward and reverse primer, 1 U KAPA2G Robust DNA Polymerase (Kapa Biosystems) and 1 μL of the cDNA or Outer RACE PCR product as PCR template. Amplification was carried out with the following program: 95℃ for 3 min, 35 cycles of 95℃ for 15 s, 58℃ for 30 s and 72℃ for 30 s, and a final extension of 5 min at 72℃. In order to confirm the obtained 5’ and 3’ cDNA ends from RACE reactions, cDNA was prepared using Oligo (dT)_18_ primer with a High-Capacity cDNA Reverse Transcription Kit (ThermoFisher Scientific). Then, nested PCR primers were designed in the 5’ and 3’ UTRs to amplify the whole coding region of *TSHR*. Targeted PCR products were purified from 1% agarose gel using QIAquick Gel Extraction Kit (Qiagen) and Sanger sequenced (Eurofins Genomics) with five primers to span the entire PCR product. All primers used for RACE reactions and full-length transcript validation are listed in Supplementary Table S4.

### Characterization of inversion breakpoint

In order to identify the breakpoints of the inversion, we implemented the BreakDancer software (Chen *et al*., 2009) on 46 individual sequenced samples independently. Reads with mapping quality above 30 were retained in the analysis. BreakDancer helped to narrow down the range of the potential breakpoints into 5 kb around the ends of the inversion. In an attempt to find the breakpoints at single base level, we extracted soft clipped reads and compared the normalized depths between samples carrying the Southern and Northern haplotypes around such reads. The candidate breakpoints are expected to occur on where the differentiation of the depth is maximized. Besides, we checked the distribution of chimeric reads across the genome. The visualization of short reads and clipped reads was performed in IGV (Robinson *et al*., 2017).

### Identification of conserved synteny across teleosts

Whole genome alignments were performed using Satsuma Chromosemble (Grabherr *et al*., 2010), and the detection of blocks of co-linearity was made with a custom R (R Core Team, 2015) script. The following genome versions were used: northern pike (*Esox lucius*): GCA_000721915.3 (Rondeau *et al*., 2014); threespine stickleback (*Gasterosteus aculeatus*): Gac-HiC_revised_genome_assembly (Peichel *et al*., 2017); guppy (*Poecilia reticulata*): GCA_000633615.2 (Kunstner *et al*., 2016); zebrafish (*Danio rerio*): GCA_000002035.4; medaka (*Oryzias latipes*):GCA_002234695.1 (Ichikawa *et al*., 2017), human (*Homo sapiens*): GRCh38; Chicken (*Gallus gallus*): GRCg6a. The zebrafish, human and chicken genomes are all from the Genome Reference Consortium (Church *et al*., 2011).

### Data Access

The assembly and RNA reads for annotations generated in this study have been submitted to the European Nucleotide Archive under accession number PRJEB31270.

## Acknowledgements.

This work was supported by The Knut and Alice Wallenberg Foundation, The Swedish Research Council and the Norwegian Research Council project 254774 (GENSINC). We thank Carl-Johan Rubin and Kerstin Howe for valuable advise during the preparation of this assembly and all Junior High School students that contributed to the project Forskarhjälpen.

## Author contributions

LA designed the study. MEP built the hybrid assembly and performed analysis. IB constructed the FALCON-assembly. GF, XH, QX, HZ, SL and XL generated the Hi-C dataset. AF cultivated fish and provided samples for linkage mapping. CHM built the linkage map. FH generated the recombination profile and breakpoint estimation. JH performed GO analysis. JC refined the *THSR* gene model. OW assisted in assembly construction and performed experimental work. LH, TH, FJM and PF performed annotation. MEP and LA wrote the manuscript with input from others. All authors approved the final version of the manuscript.

## Supplementary Material

**Supplementary Figure S1**: Linkage map and recombination-rate profile (dots) estimated using LD-hat for chromosome 1-26. Solid-line: sex-average linkage map, dashed line: male linkage map, dotted line: female linkage map.

**Supplementary Figure S2**: (*a)* Observed and expected amounts of overlapping synteny blocks. The black line is the observed amount of sequence as a function of the number of overlapping species. The red line is the expectation, if all species had the same amount of co-linear regions. (*b)* Excerpt of the ENSEMBL human genome browser (hg38) corresponding to the region on herring chromosome 6 that remains unbroken in all nine genomes.

**Supplementary Figure S3**: Sketch of repeat structures at the proposed inversion breakpoints, and read alignments for individuals of different inversion genotypes.

**Supplementary Figure S4**: (*a,b)* Histograms of allele frequency distributions in “S” (*a*) and “N” (*b*) homozygotes.

**Supplementary Table S1:** Sex-specific linkage maps.

**Supplementary Table S2:** Genes adjacent to chromosome 12 inversion breakpoints.

**Supplementary Table S3:** Sampling locations in the student project “Forskarhjälpen.”

**Supplementary Table S4:** Primers used for *TSHR* RACE.

**Supplementary Text S1:** FALCON assembly configuration file.

**Supplementary Text S2:** Annotation pipeline description.

**Supplementary Data S1:** Duplicated fragments removed from the final assembly.

